# Automated escape system: identifying prey’s kinematic and behavioral features critical for predator evasion

**DOI:** 10.1101/2023.07.02.547369

**Authors:** Nozomi Sunami, Hibiki Kimura, Hidechika Ito, Koichi Hashimoto, Yuta Sato, Soki Tachibana, Mikiya Hidaka, Kouki Miyama, Hirofumi Watanabe, Yuuki Kawabata

## Abstract

1. Identifying the kinematic and behavioral variables of prey that influence the evasion from predator attacks is essential not only for comprehending the determinants of successful predator evasion but also for shedding light on the evolution of specific traits and the dynamics of predator-prey relationships on a larger scale. However, quantifying the relationship between these variables and the success or failure of predator evasion is challenging, particularly for variables with small variations within prey species. One promising approach to address this challenge is the use of a simulated prey system, which allows us to manipulate the kinematic and behavioral features of prey and expose them to real predators. Nevertheless, creating a system that moves comparably to real prey animals remains difficult, especially for invertebrate and lower vertebrate species that respond quickly to predators and escape rapidly.
2. In this study, we have developed an automated escape system that is comparable to real prey species, responding to a predator within tens of milliseconds and escaping at over 1.0 m/s. The system automatically detects an approaching predator and pulls the prey away from the predator once the predator reaches a predetermined threshold distance. Reaction distance, response latency, as well as escaping speed, duration, and direction can be adjusted in the system.
3. By repeatedly measuring the response latency and escaping speed of the system, we demonstrated the system’s ability to exhibit fast and rapid responses while maintaining consistency across successive trials. As a case study, we manipulated the escape speed and reaction distance of the prey to expose them to a predatory fish, *Coreoperca kawamebari*. The results show that both variables significantly affect the outcome of predator-prey interactions.
4. These findings indicate that the developed escape system is useful for identifying kinematic and behavioral features of prey that are critical for predator evasion. Moreover, due to its relatively low cost and customizability, we propose that this system can be applied to investigate various aspects of animal behaviors (e.g., eliciting escape responses by artificial stimuli) in different animal species.

## Introduction

Predation is a strong selective force that shapes various forms of defensive tactics of prey (Endler 1991, Davies et al. 2012). One of the most common tactics when encountering a predator is the escape response, which includes turning swiftly and accelerating away from it (Cooper and Blumstein 2015, Domenici and Hale 2019). Numerous studies have explored the environmental and internal factors that influence the behavioral and kinematic variables associated with the escape response, such as reaction distance, escape trajectory, response latency, and speed (Cooper et al. 2003, Meager et al. 2006, Bateman and Fleming 2014, Kawabata et al. 2023). These studies assume that these variables determine the success or failure of predator evasion. However, direct evidence linking these variables to evasion outcomes is limited (Walker et al. 2005, Dangles et al. 2006, Kimura and Kawabata 2018), especially for variables with small variations within prey species. Understanding the relationship between these variables and successful predator evasion is crucial not only for comprehending the determinants of successful predator evasion but also for shedding light on the evolution of specific traits and the dynamics of predator-prey relationships on a larger scale.

One promising approach to linking the prey’s behavioral and kinematic variables with the outcome of predator-prey interactions is the use of a simulated prey system, which allows for the manipulation of the prey’s kinematic and behavioral features, exposing them to real predators. For instance, Shifferman and Eilam (Shifferman and Eilam 2004) and Ilany and Eilam (Ilany and Eilam 2008) manipulated the escape direction and flight initiation distance of prey items (i.e., dead mouse or chick) by attaching a string to the prey and manually pulling it from a distance, subjecting them to barn owl *Tyto alba* attacks. Szopa-Comley and Ioannou (Szopa-Comley and Ioannou 2022) developed a robotic prey system and manipulated the repeated escape trajectories against predatory fish, either fixed to a specific direction or randomized within a 270-degree range (45∼315 degree away from the predator). Nevertheless, creating a system that moves comparably to real prey animals remains challenging, especially for invertebrate and lower vertebrate species that respond quickly to predators (e.g., within tens of milliseconds) and escape rapidly (e.g., exceeding 1.0 m/s) (Bullock 1984).

Here, we have developed an automated escape system comparable to real prey species. The system can respond to a predator within tens of milliseconds and achieve speeds exceeding 1.0 m/s during escape. It automatically detects an approaching predator and pulls the prey away from the predator once the predator reaches a predetermined threshold distance. The system is relatively low cost (approximately 2,000 US dollars), customizable, and versatile for various applications, including eliciting escape responses in animals. This paper provides a detailed description of (1) the technical aspects of the developed system, (2) a performance test conducted to evaluate the system’s ability to exhibit fast and rapid responses while maintaining consistency across successive trials, and (3) a case study investigating effects of escape speed and reaction distance on the success or failure of evading live predatory fish.

## Materials and Methods

### Automated escape system

The system comprised a USB camera (DMK33UX287, The Imaging Source Co., Ltd., Bremen, Germany), a laptop (DAIV 5N 20075N-CML, Mouse Computer Co., Ltd., Tokyo, Japan, CPU: Intel(R) Core(TM) i7-10875H @ 2.30GHz), a microcontroller board (UNO R3, Elegoo Inc., Shenzhen, China) with a motor driver shield (SU-1201, EK Japan Co., Ltd., Fukuoka, Japan), a DC motor (RE-260RA, Mabuchi motor Co., Ltd., Chiba, Japan) with a pulley (diameter = 50 mm), and a weight connected to the pulley by a string (Figure 1). The experimental arena was surrounded by double circular plastic walls (outer: diameter = 495 mm, height = 160 mm; inner: diameter = 365 mm, height = 130 mm). These walls were attached using PVC pipes (outer: inner diameter = 30 mm, height = 35 mm; inner: inner diameter = 15 mm, height = 70 mm) with the inner wall and PVC pipes floating 15 mm above to hold the path of the weight. Two of the PVC pipes provided space for the string and weight, and had a pulley (diameter = 10 mm) attached to their lower ends to reduce friction when moving the string. The string through the PVC pipes was brought together behind the outer wall and connected to the motor with the pulley. The string consisted of a PE line (length = 2.2 m) and a transparent carbon nylon line (length = 10 m). We used a fishing barrel swivel as the weight to prevent it from rolling due to a twisted string. The escape direction of prey could be adjusted by using a different PVC pipe to pass through a string toward the motor.

**Figure 1.**
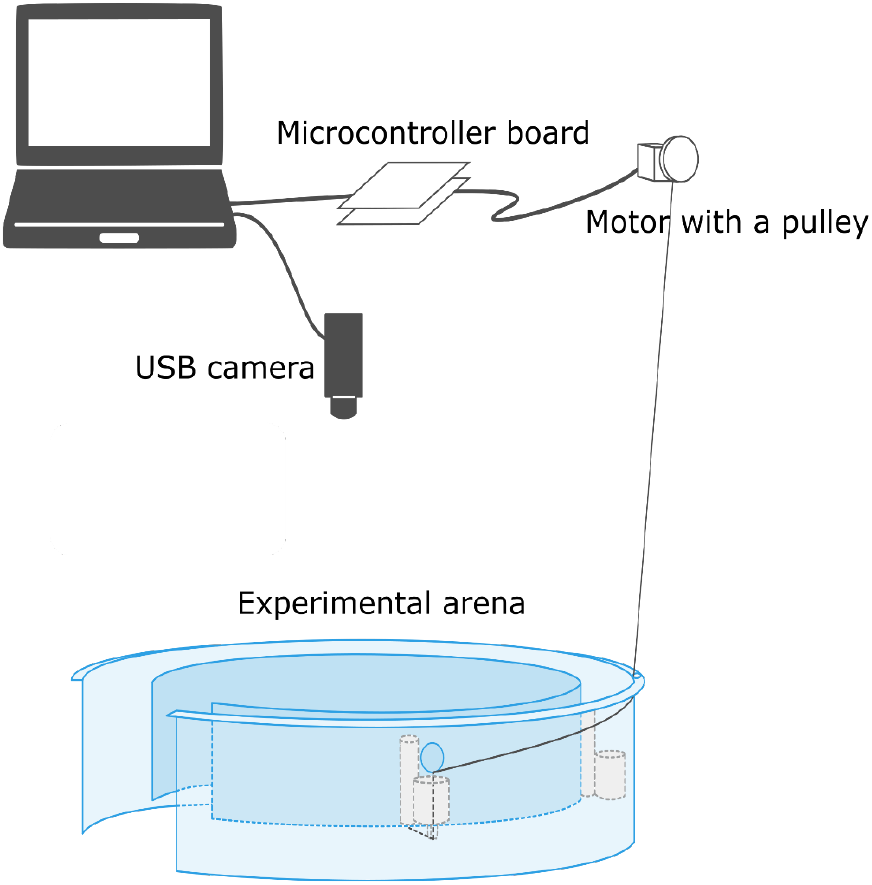
**Schematic drawing of the automated escape system**.

The step-by-step process of programs for detecting an approaching predator and triggering the escape movement of the prey is shown in Figure 2. The USB camera was installed above the experimental arena to monitor the real-time movement of the predator and display the image on the laptop. Two circles: the search circle (outer circle) and reaction circle (inner circle), were generated using a program superimposed onto the image (Figure 3). Inside the search circle, the uppermost point of the predator was detected and tracked (Figure 3). When the uppermost point of the predator was detected within the reaction circle, the laptop computer sent a signal to the microcontroller board. The custom program for the predator monitoring system was written in Python (version 3.7.7) with the OpenCV library (Opencv-contrib 3.4.10.37). In short, images were cropped to enhance their clarity, and then subjected to color thresholding for blob analysis. From the binary blob that represented the predator’s shape, its uppermost point was detected and tracked. The radius of search and reaction circles could be adjusted in the Python program. The temporal resolution (i.e., frame rate) of the USB camera was limited by the program and exhibited fluctuations with a mean±standard deviation of 238±39 fps. The custom Python program is available at https://github.com/YuukiKawabata-Lab/PreyEscapeSystem

**Figure 2.**
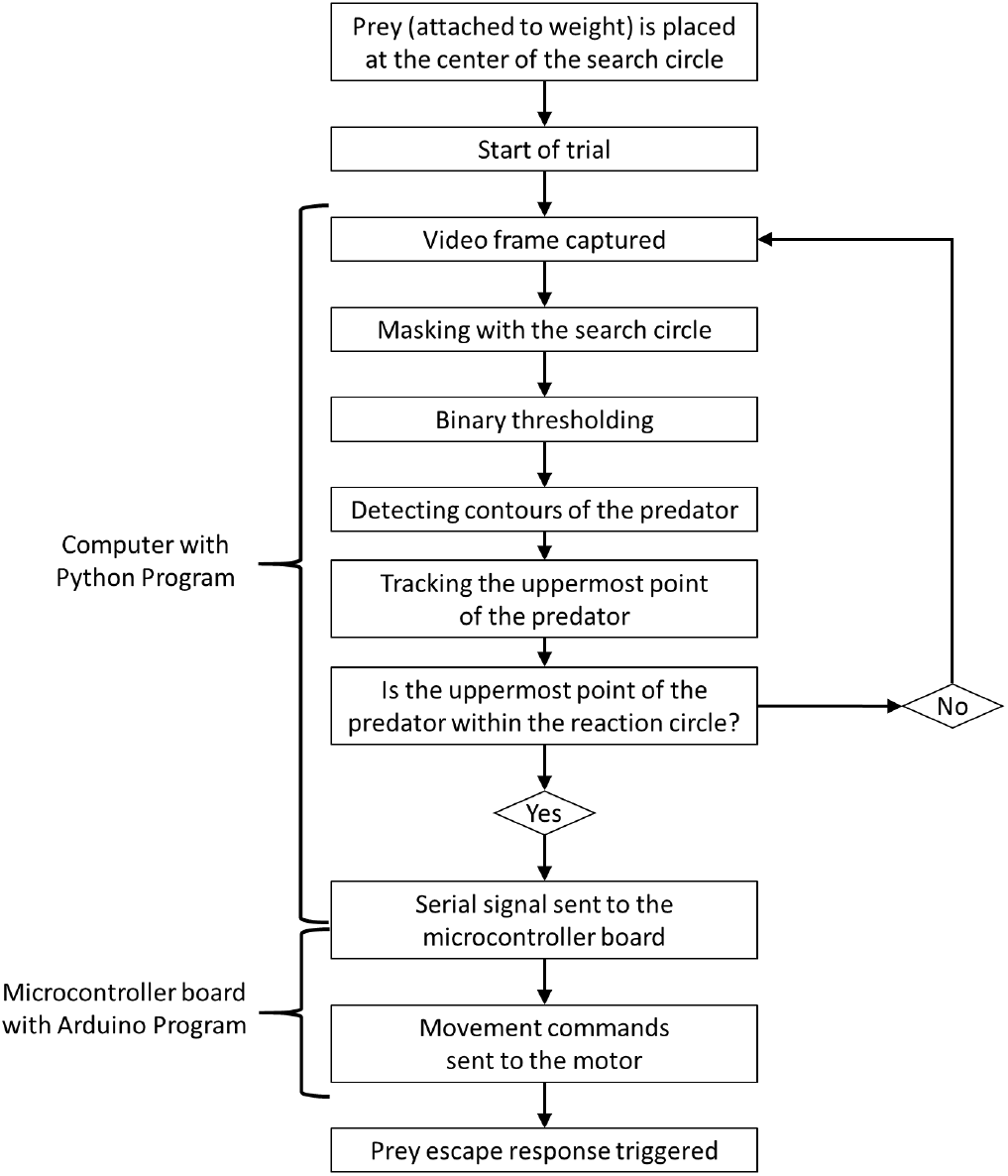
**Sequential diagram illustrating the step-by-step process of the Python and Arduino programs for detecting a predator and triggering prey escape movement**.

**Figure 3.**
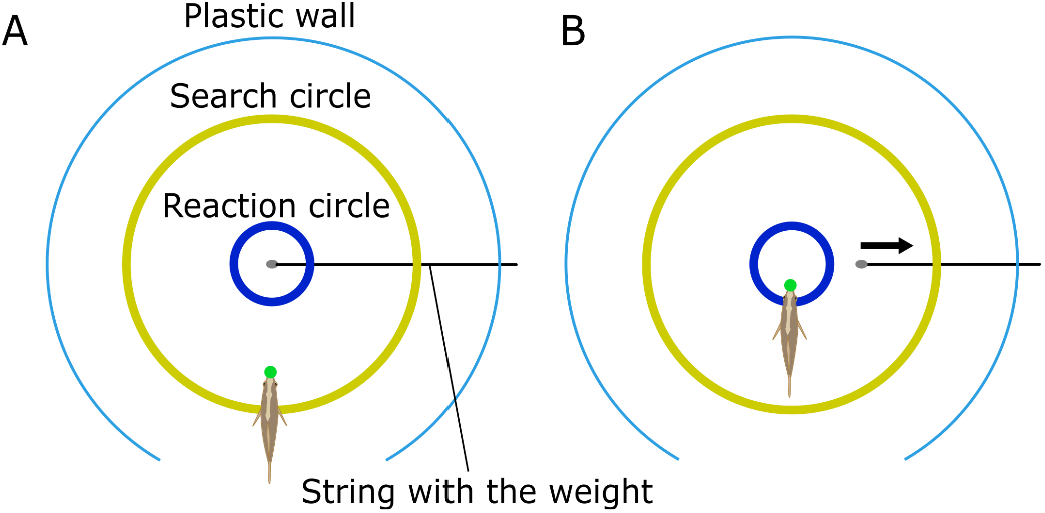
Sketch showing the top views of real-time video images with two circles overlaid. (A) When a predator is within the search circle (the outer yellow circle), its uppermost point (green-filled circle) is automatically tracked in real-time. (B) When the predator’s uppermost point is detected in the reaction circle (the blue inner circle), the laptop computer sends a signal to the microcontroller board. The board then sends movement commands to the motor, which rolls up the string. This causes the weight (with prey) attached to the string to start moving.

The movement of the prey was controlled by the motor and microcontroller board with a custom program written in Arduino IDE (version 1.8.19) through a connection with the string between the weight (the prey) and the pulley. When the microcontroller board received the predator detection signal from the laptop computer, it sent movement commands to the motor, which rolled up the string and dragged the weight (with prey) (Figure 3B). The Arduino program provided the flexibility to adjust the timing (latency) and duration of the motor’s rotation with millisecond resolution. The rotation speed of the motor could be regulated by voltage values ranging from 0 (minimum) to 255 (maximum). The custom Arduino program is available at https://github.com/YuukiKawabata-Lab/PreyEscapeSystem.

### Performance test

To evaluate the performance and consistency of the developed system, we conducted 30 trials and analyzed the response latency and moving speed of the weight. The system was placed inside a rectangular acrylic tank (width = 60 cm, height = 60 cm, length = 150 cm). Throughout the experiment, the water depth in the tank was maintained at 100 mm. The radius of the search and reaction circles were set to 153 mm and 40 mm, respectively, with the weight positioned at the center of these circles. Three different speeds (high, middle, and low) were utilized in the motor control program, corresponding to voltage values of 245 (high), 40 (middle), and 11 (low). We conducted 10 trials at each programmed speed. In every trial, the motor rotation was programmed to displace the weight approximately 232 mm from its starting position towards the PVC pipe on the right side. Delay time was not included in the program as our objective was to estimate the baseline response latency of the system. The system was activated by manually moving a dummy predator into the inner circle. The movements of the weight were recorded from above using a high-speed video camera (RX10IV, Sony Corp., Tokyo, Japan) with a frame rate of 480 fps.

The recorded videos were analyzed using Kinovea ver. 0.8.27 (www.kinovea.org). The response latency was determined as the duration between the entry of the dummy predator into the reaction circle and the initiation of weight movement. To determine the moving speed, the coordinates of the weight were digitized in each frame from its starting position to the inner plastic wall. Subsequently, we calculated the time-series speed of the weight from its positional data using Lancoz smoothing method. The effect of three different programmed speeds on response latency was examined using one-way analysis of variance (ANOVA). In order to assess the consistency of the measurements, we calculated the coefficients of variation (CVs) for the response latency, as well as the mean and maximum speeds corresponding to each programmed speed. Consistency of the parameters were classified based on the previous study (Aronhime et al. 2014), defining excellent when CV was <10%, good when CV was 10∼20%, acceptable when CV was 20∼30%, and poor when CV was >30%. All analyses were carried out using R (version 4.0.5).

### Case study

To demonstrate the utility of the system, we exposed the controlled prey to a live predatory fish *C. kawamebari* (total length: 69.2 mm). We manipulated the escape speed (i.e., mean speed from the starting position to the inner plastic wall; Figure 2) and reaction distance (i.e., radius of the reaction circle; Figure 2) to investigate whether these variables influence the predation probability. This experiment was conducted using the same tank and system in the performance test. The fish had been kept in a compartment (width = 20 cm, height = 25 cm, length = 20 cm) made with transparent PVC punching plates. Live shrimp *Neocaridina denticulata* (total length: 14.4±1.4 mm) attached to the weight with adhesive was used as prey. Live prey was chosen because the preliminary experiment showed that the predator did not chase or strike towards dead prey or artificial pellets. The prey’s heads were positioned to face the entrance of the experimental arena. At the start of each trial, the compartment was positioned in front of the arena’s entrance, laid horizontally to allow the fish to enter the arena. As in the performance test, the radius of the search circle was set to 153 mm, and the system was programmed to move the weight 232 mm from its starting position towards the PVC pipe on the right side. A total of 33 trials were conducted, with the prey being randomly assigned seven different speeds and eight different reaction distances to elicit the predator’s movement. Because the actual escape speed cannot be measured when the predator successfully captures the prey, we measured the speed of the actual movements five times for each programmed speed before the trial (without the predator), and its mean value was used for the analysis. The water depth was kept at 100 mm and temperature was maintained at 22.9±0.2 °C throughout the experiment.

In a few cases, the predator grabbed or touched the prey, but the predator finally failed to capture the prey. Because this study focused on kinematic and behavioral variables rather than the other variables (e.g. size, spines), these cases were regarded as successful capture. The effects of prey’s escape speed and reaction distance on predation probability were evaluated using a logistic regression analysis. Success and failure of predation were designated as 1 and 0, respectively, and used as the objective variable. Escape speed and reaction distance were considered as the explanatory variables.

The significance of the variables was assessed by removing them from the model, and comparing the change in deviance using the likelihood ratio test with a χ^2^ distribution. The analysis was carried out using R (version 4.0.5).

## Results

### Performance test

There was no significant difference in the response latency among the three programmed speeds (Table 1; ANOVA, *F*_(2, 27)_=0.82, *P*=0.45). The mean response latency was 72.0±7.4 ms (mean±standard deviation), indicating that the weight started moving about 72 ms after the dummy predator entered the reaction circle (Table 1; Figure 4). This value is comparable to the response latencies of visually-evoked escape responses in different fish species (Batty 1989, Paglianti and Domenici 2006, Dunn et al. 2016, Cade et al. 2020) (Figure 4). With a CV of the response latency at 10.2%, the consistency of the response latency in this system is considered good and nearly excellent.

**Table 1.** Summary of the performance test.

| Programmed speed | Response Latency (ms) | Mean Speed (m/s) | Maximum Speed (m/s) |
| --- | --- | --- | --- |
| High | 73.7±8.6 (11.7%) | 1.83±0.04 (2.0%) | 2.65±0.11 (4.0%) |
| Middle | 72.6±7.9 (10.8%) | 0.89±0.02 (1.8%) | 1.23±0.06 (4.5%) |
| Low | 69.6±5.3 (7.7%) | 0.29±0.01 (1.7%) | 0.41±0.03 (8.2%) |
| Total | 72.0±7.4 (10.2%) | - | - |

Mean±standard deviation of the value is shown. The coefficient of variation is shown in the parenthesis.

**Figure 4.**
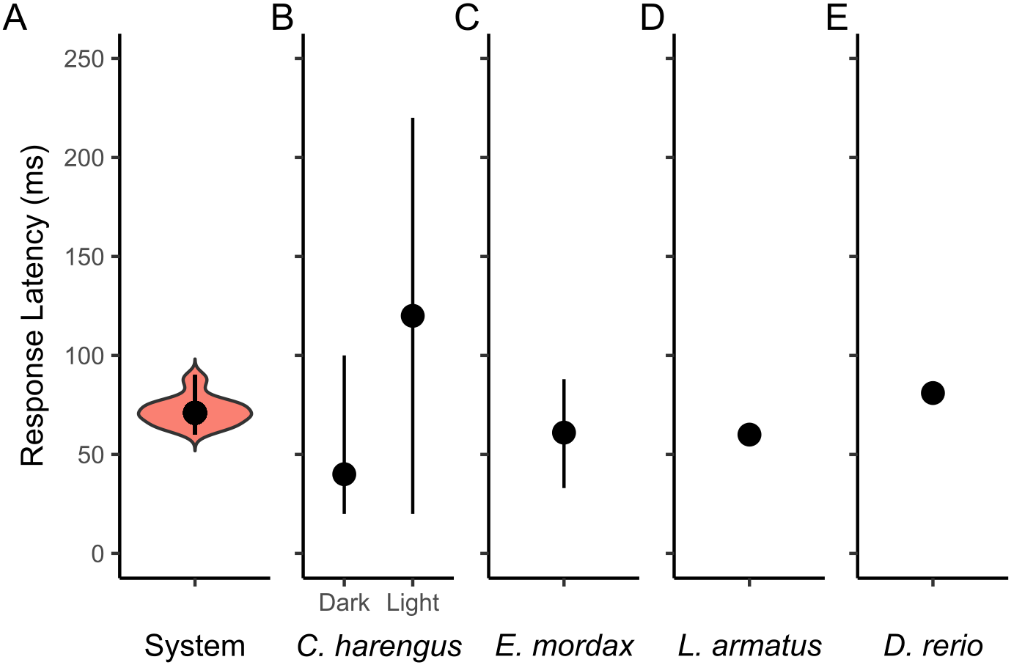
The response latency of the developed system and those of the visually-evoked escape responses in different fish species. The black filled dot and vertical line represent the median and range of the values, respectively. (A) Response latency of the developed system. The width of the red filled area (violin plot) represents the kernel probability density. (B) Response latency of larval herring *Clupea harengus* to a flush light under dark and light conditions. Data were obtained from Figure 4 in (Batty 1989). (C) Response latency of anchovy *Engraulis mordax* to a flush light. Data were obtained from (Cade et al. 2020). (D) Response latency of staghorn sculpin *Leptocottus armatus*, estimated from the fish’s latest response after the end of the expansion of the looming stimulus (Paglianti and Domenici 2006). (E) Response latency of larval zebrafish *Danio rerio*, estimated from the escape responses to looming stimuli with different expansion speeds (Dunn et al. 2016).

Time-series moving speeds of the weight (prey) for three different programmed speeds are shown in Figure 5. The variations among trials were apparently small, and the CVs of the mean and maximum speeds for all three programmed speeds were less than 10% (Table 1). This suggests that the consistency of the moving speeds of the system is excellent. When the speed was set to low, the weight rapidly accelerated to about 0.3 m/s in around 5 ms and maintained that speed. When the speed was set to middle or high, the weight reached the inner wall before its moving speed stabilized. For the middle speed set, the weight rapidly accelerated to approximately 0.7 m/s in around 25 ms, followed by gradual acceleration with oscillations. For the high speed set, the weight rapidly accelerated to approximately 1.8 m/s in about 15 ms, and then gradually accelerated to about 2.5 m/s by the 40 ms mark. The speed then decelerates to 1.5 m/s and eventually reached the inner plastic wall.

**Figure 5.**
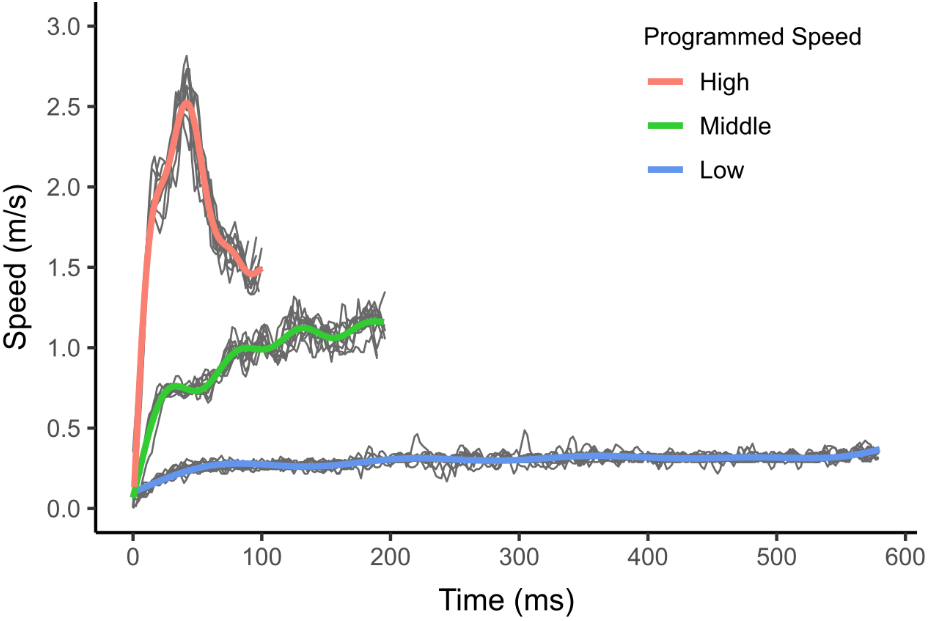
Time-series moving speeds of the weight (prey) in the developed escape system. Three different speeds (high, middle, and low) were utilized in the motor control program, and 10 trials were conducted at each programmed speed. The thin gray line represents the moving speed of each trial, while the thick colored line represents the smoothed curve of each programmed speed, estimated by the generalized additive mixed model.

**Figure 6.**
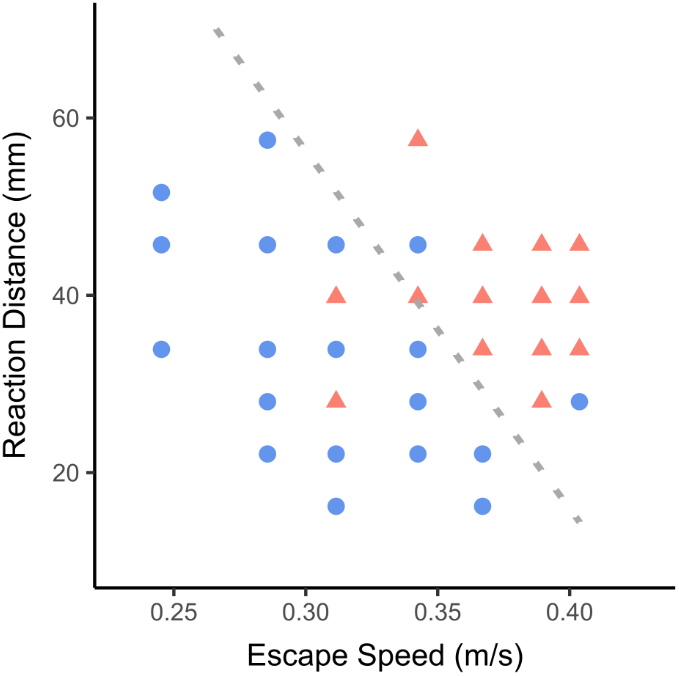
Effects of escape speed and reaction distance of prey (manipulated by the developed escape system) on the outcome of predator-prey interactions. Blue circles and red triangles are indicative of capture and escape, respectively. The dotted line represents the 50% predation probability estimated from the logistic regression analysis, with the area below and above the line indicating predicted capture and escape, respectively. Out of the 33 data points analyzed, 29 (87.9%) were correctly categorized by the estimated line.

### Case study

The success of predation was found to be influenced by both escape speed and reaction distance (Figure 5; LR test, escape speed: χ^2^=17.39, d.f.=1, *P*<0.01; reaction distance: χ^2^=5.70, d.f.=1, *P*<0.05). Faster escape speeds and longer reaction distances were associated with a lower likelihood of predation success. The odds ratio, based on one standard deviation, was 0.64 for escape speed and 0.33 for reaction distance. These findings are consistent with previous studies that used live prey and predators (Walker et al. 2005, Langerhans 2009, Stewart et al. 2013, Kimura and Kawabata 2018).

## Discussion

The performance test confirms that the developed escape system responds quickly to an approaching predator and escapes rapidly. Furthermore, it exhibits consistent performance across successive trials, ensuring reproducibility. The case study involving a live predator reveals that the manipulated parameters (i.e., escape speed and reaction distance) significantly affect the success of evading predator attacks, consistent with the findings of previous studies that used live prey and predators (Walker et al. 2005, Langerhans 2009, Stewart et al. 2013, Kimura and Kawabata 2018). These findings indicate that the developed system is useful for identifying the prey’s kinematic and behavioral features that are crucial for predator evasion.

When the prey was programmed to move at middle and high speed, oscillations were observed in the time-series speed of prey movement (Figure 5). These oscillations can be attributed to the extension and contraction of the string. According to the principles of physics, a force exceeding the static friction force is required to initiate the weight movement. Therefore, the string attached to the weight would have been slightly extended before the onset of the weight movement, which caused rapid accelerations after the onset. The rapid accelerations would then contract the string, resulting in decelerations of the weight movement. This process is likely to have produced the oscillations in the speed of prey movement. Using an inextensible string and/or a weight made with a material that has less friction can help alleviate the oscillations in the prey’s speed over time. Nonetheless, it is worth noting that these kinds of oscillation patterns are often observed in actual animal movements (Alexander 2003, Voesenek et al. 2019).

Previous studies have used different simulated prey systems that manipulate the kinematic and/or behavioral variables of prey to study their interactions with live predators. The simplest method involves attaching a string to the prey and manually pulling it from a distance (Shifferman and Eilam 2004, Ilany and Eilam 2008). While this method is easy to implement, it is limited by the response latency of human visual processing, which is approximately 175∼202 ms (Solanki et al. 2012). Additionally, the escape speed of the prey is influenced by human performance, which can lead inconsistencies across successive trials. A virtual prey system, in which prey objects are projected onto one side of an experimental tank, has been used to examine the schooling parameters of prey by manipulating their movement trajectories (Ioannou et al. 2012, Duffield and Ioannou 2017, Ioannou et al. 2019). However, live predators are generally attracted to the movement of virtual prey, and virtual prey is not programmed to respond to approaching predators. Therefore, this method is not suitable for situations where the prey remains stationary at first and rapidly initiates escape motion. Additionally, because predators are not rewarded for successful attacks on virtual prey, this system cannot address questions that require successive trials for individual predators to learn (e.g., random vs specific escape trajectories). The recently developed robotic prey system (Swain et al. 2012, Szopa-Comley and Ioannou 2022), which responds to approaching predators by turning and escaping, can be successfully used to answer such questions. However, it has a longer response latency (approximately 700 ms) likely due to its complex system. In addition, its achievable maximum speed may be lower than that of our system due to the robotic prey’s heavier mass. Consequently, it may be challenging for the robotic prey system to respond quickly and escape rapidly to evade actual predators, making it difficult to directly link the prey’s variables to successful predator evasion. In the previous study (Szopa-Comley and Ioannou 2022), indirect metrics (i.e., time required for the predator to capture the prey) was used to examine the benefit of random escape directions over specific directions. In contrast, our system can control the prey’s escape speed over a relatively wide range, including speeds that enable prey to evade actual predator attacks. Additionally, it reacts to approaching predators as quickly as real prey species (i.e., <100 ms). Therefore, our developed system is more suitable for certain research questions and complements previous methods, including virtual and robotic prey systems. While our current system is not designed for multiple individuals, the number of prey and escape direction choices can be easily increased by using multiple strings and motors, enabling simulations of grouped prey escaping in various directions.

One potential drawback of the developed system is that the prey’s escape direction may be predictable to predators due to the presence of the string. This becomes problematic, especially when the research question involves examining escape directions across successive trials (e.g., random vs specific escape directions) and predators can perceive or sense the string. To address this issue, one possible solution is to incorporate a magnet system similar to the one used in the robotic prey system (Swain et al. 2012, Szopa-Comley and Ioannou 2022). In this setup, one magnet connected to a motor is positioned underneath the floor of the experimental arena, while another magnet is attached to the prey on the floor. However, it is important to note that using a magnet may increase friction, and result in slower prey movement and larger oscillation amplitudes in speed. Another potential solution is to use multiple strings placed underneath the weight, which could prevent predators from predicting the escape direction. Further research is necessary to investigate whether predators can perceive transparent strings and to evaluate the effectiveness of modified systems, such as those utilizing magnets or multiple strings.

Our system has potential to be applied to a range of objectives beyond predator evasion. For instance, it can be utilized to investigate the spatial learning capabilities of animals by modifying the shape of the reaction circle to create blind zones for the prey. This approach enables the examination of whether and how animals learn specific spatial rules and adapt their behavior accordingly. Another potential application is the elicitation of escape responses in prey species. Many studies have employed artificial stimuli such as dummy predators, air-puffs, or dropping balls, to trigger escape responses in prey (Camhi and Tom 1978, Meager et al. 2006, Marras and Domenici 2013, Kawabata et al. 2023). In these studies, observers typically wait for the prey to approach a specific location or force the prey to stay at that location before triggering the artificial stimulus. Our system can be modified to efficiently collect data in such cases by placing the reaction circle in front of the artificial stimulus and connecting the microcontroller board to the device that generates the artificial stimulus. These examples demonstrate the versatility of our device, which can be utilized for investigating various questions in animal behavior beyond predator evasion.

In this study, we have introduced a novel prey escape system that demonstrates fast and rapid responses to predators while maintaining consistency across successive trials. Our system allows for the manipulation of various kinematic and behavioral variables, including reaction distance, response latency, escaping speed, duration, and direction. This feature enables us to establish a direct link between these prey variables and successful predator evasion. Furthermore, the relatively low cost and customizability of our system make it a versatile tool that can be applied to investigate diverse aspects of animal behavior in different species. By adapting and implementing this system, researchers can delve into a wide range of research questions related to predator-prey dynamics and explore other fascinating behaviors exhibited by animals. Overall, our developed prey escape system holds great potential for advancing our understanding of animal behavior and enhancing our knowledge of predator-prey interactions in various ecological contexts.

## Acknowledgements

The animal care and experimental procedures were approved by the Animal Care and Use Committee of the Faculty of Fisheries (Permit No. NF-0058), Nagasaki University in accordance with the Guidelines for Animal Experimentation of the Faculty of Fisheries and the Regulations of the Animal Care and Use Committee, Nagasaki University. This study was funded by Grants-in-Aid for Scientific Research, Japan Society for the Promotion of Science, to Y.K. (19H04936).

## References

Alexander, R. M. 2003. Principles of animal locomotion. Princeton University Press, New Jersey.

Aronhime, S., C. Calcagno, G. H. Jajamovich, H. A. Dyvorne, P. Robson, D. Dieterich, M. I. Fiel, V. Martel-Laferriere, M. Chatterji, H. Rusinek, and B. Taouli. 2014. DCE-MRI of the liver: effect of linear and nonlinear conversions on hepatic perfusion quantification and reproducibility. Journal of Magnetic Resonance Imaging 40:90–98.

Bateman, P. W., and P. A. Fleming. 2014. Switching to Plan B: Changes in the escape tactics of two grasshopper species (Acrididae: Orthoptera) in response to repeated predatory approaches. Behavioral Ecology and Sociobiology 68:457–465.

Batty, R. S. 1989. Escape responses of herring larvae to visual stimuli. Journal of the Marine Biological Association of the United Kingdom 69:647–654.

Bullock, T. H. 1984. Comparative neuroethology of startle, rapid escape, and giant fiber-mediated responses. Pages 1-13 in R. C. Eaton, editor. Neural mechanisms of startle behavior. Springer, Boston, MA.

Cade, D. E., N. Carey, P. Domenici, J. Potvin, and J. A. Goldbogen. 2020. Predator-informed looming stimulus experiments reveal how large filter feeding whales capture highly maneuverable forage fish. Proc Natl Acad Sci U S A 117:472–478.

Camhi, J. M., and W. Tom. 1978. The escape behavior of the cockroach Periplaneta americana - I. Turning response to wind puffs. Journal of Comparative Physiology A 128:193–201.

Cooper, W. E., and D. T. Blumstein. 2015. Escaping from predators. Cambridge University Press, Cambridge, UK.

Cooper, W. E., V. Pérez-Mellado, T. Baird, T. A. Baird, J. P. Caldwell, and L. J. Vitt. 2003. Effects of risk, cost, and their interaction on optimal escape by nonrefuging Bonaire whiptail lizards, Cnemidophorus murinus. Behavioral Ecology 14:288–293.

Dangles, O., N. Ory, T. Steinmann, J. P. Christides, and J. Casas. 2006. Spider’s attack versus cricket’s escape: velocity modes determine success. Animal Behaviour 72:603–610.

Davies, N. B., J. R. Krebs, and S. A. West. 2012. An introduction to behavioural ecology. Fourth edition. Wiley-Blackwell, Oxford, UK.

Domenici, P., and M. E. Hale. 2019. Escape responses of fish: a review of the diversity in motor control, kinematics and behaviour. Journal of Experimental Biology 222.

Duffield, C., and C. C. Ioannou. 2017. Marginal predation: do encounter or confusion effects explain the targeting of prey group edges? Behav Ecol 28:1283–1292.

Dunn, T. W., C. Gebhardt, E. A. Naumann, C. Riegler, M. B. Ahrens, F. Engert, and F. Del Bene. 2016. Neural circuits underlying visually evoked escapes in larval zebrafish. Neuron 89:613–628.

Endler, J. A. 1991. Interactions between predators and prey. Pages 169–196 in J. R. Krebs and D. N. B., editors. Behavioural Ecology: an Evolutionary Approach. Blackwell Scientific Publications, Oxford, UK.

Ilany, A., and D. Eilam. 2008. Wait before running for your life: defensive tactics of spiny mice (Acomys cahirinus) in evading barn owl (Tyto alba) attack. Behavioral Ecology and Sociobiology 62:923–933.

Ioannou, C. C., V. Guttal, and I. D. Couzin. 2012. Predatory fish select for coordinated collective motion in virtual prey. Science 337:1212–1215.

Ioannou, C. C., F. Rocque, J. E. Herbert-Read, C. Duffield, and J. A. Firth. 2019. Predators attacking virtual prey reveal the costs and benefits of leadership. Proc Natl Acad Sci U S A 116:8925–8930.

Kawabata, Y., H. Akada, K. I. Shimatani, G. N. Nishihara, H. Kimura, N. Nishiumi, and P. Domenici. 2023. Multiple preferred escape trajectories are explained by a geometric model incorporating prey’s turn and predator attack endpoint. eLife 12.

Kimura, H., and Y. Kawabata. 2018. Effect of initial body orientation on escape probability of prey fish escaping from predators. Biology Open 7:bio023812.

Langerhans, R. B. 2009. Morphology, performance, fitness: functional insight into a post-Pleistocene radiation of mosquitofish. Biol Lett 5:488–491.

Marras, S., and P. Domenici. 2013. Schooling fish under attack are not all equal: some lead, others follow. PLoS ONE 8.

Meager, J. J., P. Domenici, A. Shingles, and A. C. Utne-Palm. 2006. Escape responses in juvenile Atlantic cod Gadus morhua L.: the effects of turbidity and predator speed. Journal of Experimental Biology 209:4174–4184.

Paglianti, A., and P. Domenici. 2006. The effect of size on the timing of visually mediated escape behaviour in staghorn sculpin Leptocottus armatus. Journal of Fish Biology 68:1177–1191.

Shifferman, E., and D. Eilam. 2004. Movement and direction of movement of a simulated prey affect the success rate in barn owl Tyto alba attack. Journal of Avian Biology 35:111–116.

Solanki, J., N. Joshi, C. Shah, M. Hb, and G. Pa. 2012. A study of correlation between auditory and visual reaction time in healthy adults. International Journal of Medicine and Public Health 2:36–38.

Stewart, W. J., G. S. Cardenas, and M. J. McHenry. 2013. Zebrafish larvae evade predators by sensing water flow. Journal of Experimental Biology 216:388–398.

Swain, D. T., I. D. Couzin, and N. Ehrich Leonard. 2012. Real-time feedback-controlled robotic fish for behavioral experiments with fish schools. Proceedings of the IEEE 100:150–163.

Szopa-Comley, A. W., and C. C. Ioannou. 2022. Responsive robotic prey reveal how predators adapt to predictability in escape tactics. Proc Natl Acad Sci U S A 119:e2117858119.

Voesenek, C. J., R. P. M. Pieters, F. T. Muijres, and J. L. van Leeuwen. 2019. Reorientation and propulsion in fast-starting zebrafish larvae: an inverse dynamics analysis. Journal of Experimental Biology 222.

Walker, J. A., C. K. Ghalambor, O. L. Griset, D. McKenney, and D. N. Reznick. 2005. Do faster starts increase the probability of evading predators? Functional Ecology 19:808–815.

